# Assessment of lisavanbulin (BAL101553) as an anti-lymphoma agent

**DOI:** 10.1101/2022.12.31.521730

**Authors:** Filippo Spriano, Luca Aresu, Luciano Cascione, Giorgia Risi, Alberto J. Arribas, Sara Napoli, Andrea Rinaldi, Nicole Forster-Gross, Felix Bachmann, Marc Engelhardt, Heidi Lane, Francesco Bertoni

## Abstract

Microtubules are major components of the cellular cytoskeleton, ubiquitously founded in all eukaryotic cells. They are involved in mitosis, cell motility, intracellular protein and organelle transport, and maintenance of cytoskeletal shape. Avanbulin (BAL27862) is a microtubule-targeted agent (MTA) that promotes tumor cell death by destabilization of microtubules. Due to its unique binding to the colchicine site of tubulin, differently from other MTAs, avanbulin has previously shown activity in solid tumor cell lines. Its prodrug, lisavanbulin (BAL101553), has shown early signs of clinical activity, especially in tumors with high EB1 expression. Here, we assessed the preclinical anti-tumor activity of avanbulin in diffuse large B cell lymphoma (DLBCL) and the pattern of expression of EB1 in DLBCL cell lines and clinical specimens.

Avanbulin showed a potent *in vitro* anti-lymphoma activity, which was mainly cytotoxic with potent and rapid apoptosis induction. Median IC50 was around 10nM in both activated B cell like (ABC) and germinal center B cell like (GCB) DLBCL. Half of the cell lines tested showed an induction of apoptosis already in the first 24h of treatment, the other half in the first 48h. EB1 showed expression in DLBCL clinical specimens, opening the possibility for a cohort of patients that could potentially benefit from treatment with lisavanbulin.

These data show the basis for further preclinical and clinical evaluation of lisavanbulin in the lymphoma field.

## Introduction

Microtubules are major components of the cellular cytoskeleton. They are ubiquitously found in all eukaryotic cells being involved in mitotic activity, cell motility, intracellular protein and organelle transport, and maintenance of cytoskeletal shape ^1^. Microtubules are formed by the polymerization of tubulin into hollow tubes of about 250 Å in diameter. Tubulin in turn is a dimer consisting of α-tubulin and β-tubulin subunits ^2,3^. Tubulin polymerization occurs by a nucleation–elongation mechanism, in which the slow formation of a short microtubule “nucleus” is followed by rapid elongation of the microtubule at its ends by the reversible addition of α- and β-tubulin heterodimers ^4^. Microtubules assembly dynamics are a well-regulated process through anchorage on microtubule organizing centers such as centrosomes, and through interaction with a variety of proteins. Microtubules assembly and disassembly occur simultaneously and the net effect, elongation versus shortening, is determined by several factors, including the concentration of free tubulin, chemical mediators such as magnesium and calcium ions and several proteins that interact with unpolymerized tubulin and/or microtubules. Due to their central role in cellular processes, including cellular division, microtubules still represent one of the main targets for anti-cancer therapy ^5–7^. In the lymphoma field, successful examples of microtubule-targeted agents (MTAs) are vincristine and vinblastine, part of the R-CHOP and ABVD regimens, respectively, and monomethylauristatin E (MMAE), which is the payload in both brentuximab vedotin and polatuzumab vedotin ^5,8^.

Avanbulin (BAL27862) is an MTA characterized by the ability to bind to the colchicine site of tubulin, differently from other MTAs ^9^. Avanbulin is active in various preclinical models of solid tumors, including in cells resistant to drugs that bind to either the taxane or the vinca alkaloid site on tubulin or that are substrates of the multidrug resistance pump MDR1 ^10–14^. Its highly soluble prodrug, lisavanbulin (BAL101553), ^15^ has entered the initial clinical evaluation with early signs of clinical activity in patients with advanced solid tumors and with recurrent glioblastoma ^16–18^. Clinical and laboratory data suggest that the anti-tumor activity of avanbulin and of its prodrug lisavanbulin is higher in tumors with strong expression of end-binding protein 1 (EB1) ^18–20^, a microtubule-associated protein, important for the regulation of microtubule dynamics ^21^.

Based on the sensitivity of lymphomas to MTAs, here, we evaluated the *in vitro* activity of avanbulin in diffuse large B cell lymphoma (DLBCL) cell lines and its relationship with expression levels of EB1. The latter was also more extensively studied across datasets of lymphoma clinical specimens.

## Methods

### Assessment of anti-lymphoma activity of avanbulin in a panel of lymphoma cell lines

Lymphoma cell lines derived from activated B cell like (ABC) and germinal center B cell like (GCB) DLBCL were exposed to increasing doses of avanbulin. Antiproliferative activity was assessed as previously described ^22,23^. Briefly, cells were manually seeded in 96-well plates at a concentration of 50,000 cells/mL (10,000 cells in each well). Treatments were performed by using the Tecan D300e Digital Dispenser (Tecan, Mannedorf, Switzerland). After 72 hours, cell viability was determined by using 3-(4,5-dimethylth-iazol-2-yl)-2,5-dimethyltetrazolium bromide and the reaction stopped after 4 hours with sodium dodecyl sulfate lysis buffer. The identity of DLBCL cell lines was validated by short tandem repeat DNA fingerprinting ^24^.

Apoptosis induction and cell proliferation under treatment was evaluated with annexin V staining (AV+) on cells incubated in a live cell imaging system (Incucyte). Ninety-six-well plates coated with 0.01% poly-L-ornithine solution for 1 hour RT and then dried for 30 minutes prior to cell addition. 40’000 cells per well, in a total volume of 100 μL, were seeded. Cells were then treated with four different concentrations of avanbulin (5, 10, 20, 40 nM) or DMSO. Finally, annexin V reagent was added (Incucyte Annexin V Green Dye for Apoptosis) to monitor the apoptosis induction. Each condition was done in triplicate and for each well nine images were taken. Plates were then leaved into the Incucyte Live-Cell Analysis System to monitor apoptosis and cell growth. Images were taken every four hours for a total of 72 hours monitoring. The analysis was performed, with the Incucyte software, in terms of cell growth (cell-by-cell analysis) that allows the counting of the exact number of cells in each image and in terms of apoptosis, counting the number of cells GFP positive during time. Cell growth and number of GFP positive cells was normalized on time 0.

Cell cycle analysis was performed after 24, 48 and 72h of drug exposure at 20nM or DMSO. Cells were fixed and permeabilized with ethanol and subsequently stained with propidium iodide. The analyses were performed with the FlowJo software and percentages of cells in G1, S and G2/M phases of the cell cycle were determined.

### Transcriptome profiling of lymphoma cell lines

For each cell line, 5×10^6^ of cells in exponential growth were collected and resuspended in 1mL of TRI Reagent (Sigma Aldrich, Buchs, Switzerland) for cell lysis. Extraction was performed by MonarchTotal RNA Miniprep kit (New England Biolabs, Ipswich, MA, USA) according to manufacturer extraction to separate RNA molecules shorter and longer than 200 nucleotides in two different fractions. Genomic DNA was digested at the initial step of extraction. Only RNA longer than 200 nucleotides was used for library preparation, after quality check by Agilent BioAnalyzer (Agilent Technologies, Santa Clara, CA, USA) using the RNA 6000 Nano kit (Agilent Technologies) and concentration was determined by the Invitrogen Qubit (Thermo Fisher Scientific) using the RNA BR reagents (ThermoFisher Scientific, Waltham, MA, USA). The TruSeq RNA Sample Prep Kit v2 for Illumina (Illumina, San Diego, CA, USA) was used for cDNA synthesis and addition of barcode sequences. The sequencing of the libraries was performed via a paired end run on a NextSeq500 Illumina sequencer (Illumina). At least 50×10^6^ of reads were collected per each sample. The RNA-seq reads quality was evaluated with FastQC (v0.11.5) and removed low-quality reads/bases and adaptor sequences using Trimmomatic (v0.35). The trimmed-high-quality sequencing reads were aligned using STAR, a spliced read aligner which allows for sequencing reads to span multiple exons. On average, 85% of the sequencing reads were aligned for each sample to the reference genome (HG38). The HTSeq-count software package was then used for quantification of gene expression with GENCODE v22 as gene annotation. Data were subsetted to genes that had a counts-per-million value greater than five in at least one cell line. The data were normalized using the ‘TMM’ method from the edgeR package and transformed to log2 counts-per-million using the edgeR function ‘cpm’. Expression values are available at the National Center for Biotechnology Information (NCBI) Gene Expression Omnibus (GEO; http://www.ncbi.nlm.nih.gov/geo) database under the accession number GSE221770.

### In silico assessment of EB1 in lymphoma clinical specimens and its association with MYC deregulation

We accessed in-silico the association of EB1 and MYC expression in different public gene expression datasets of gene expression. We pre-processed individual datasets with ad-hoc pipelines that use the standard de-facto algorithms for the platform used for measuring the mRNA expression levels. In details, for datasets GSE4475 ^25^ and GSE22470 ^26^ probe level normalization was done using the calibration and variance stabilization method by Huber et al. ^27^. Probe-set summarization was performed using the median polish method on the normalized data ^28^. For GSE10846 ^29^, data were analyzed with Microarray Suite version 5.0 (MAS 5.0) using Affymetrix default analysis settings and global scaling as normalization method. The trimmed mean target intensity of each array was arbitrarily set to 500. For TCGA – DLBCL dataset, the alignment step was performed using a two-pass method with STAR. Quality assessment was performed pre-alignment with FASTQC and post-alignment with Picard Tools. Following alignment, BAM files are processed through the RNA Expression Workflow to determine RNA expression levels. The reads mapped to each gene are enumerated using HT-Seq-Count. Expression values are provided in a tab-delimited format. GENCODE v22 was used for gene annotation. RNA-Seq expression level read counts produced by HT-Seq are normalized with the TMM method and log2 transformed (counts-per-million). For the dataset from Schmitz et al. ^30^, paired-end reads were mapped to the human genome (NCBI build 37) using the gapped aligner STAR using the two-pass method. The alignment file was used for calculating the raw gene expression values by HTseq-count, using the intersection-nonempty model. The counts for gene expression were normalized and log2 transformed using the trimmed-mean (90%) method. For each dataset, we calculated the Pearson correlation coefficient between the normalized expression values for EB1 and MYC.

### Immunohistochemistry of DLBCL clinical specimens

Tissue samples from lymphoma clinical specimens prepared as a tissue microarrays (TMA) (LY1001D and LY2086B, collected under HIPPA approved protocols; TissueArray.com) were stained for EB1 using a CE-marked immunohistochemistry Clinical Trial Assay (Discovery Life Sciences Biomarker Services GmbH, Kassel Germany). EB1-positivity was quantified by a board-certified pathologist based on the percentage of tumor cells showing moderate or strong cytoplasmic and membrane staining for EB1, where cellular EB1 staining was defined as: intensity 0 = EB1 staining absent; intensity 1 = weak EB1 staining; intensity 2 = moderate EB1 staining; intensity 3 = strong EB1 staining.

## Results

### Assessment of anti-lymphoma activity of avanbulin in a panel of lymphoma cell lines

The *in vitro* anti-lymphoma activity of avanbulin was assessed in 26 cell lines derived from activated B cell like (ABC; n.=7) and germinal center B cell type (GCB; n.=19) DLBCL. Antiproliferative activity was first evaluated by MTT assay after 72 hours of exposure to increasing concentrations of avanbulin. The compound showed a very high activity in all the cell lines tested, with a median IC50 across cell lines of 11 nM (95% C.I., 10.03 – 16.17) (Figure 1A). No differences were observed based on the cell of origin of the cell lines in terms of IC50 values (ABC, median IC50 10 nM; GCB, 12 nM), nor in terms of AUC (ABC, median AUC 4102, GCB, 4551) (Figure1B, C, Supplementary Table 1).

**Figure 1.**
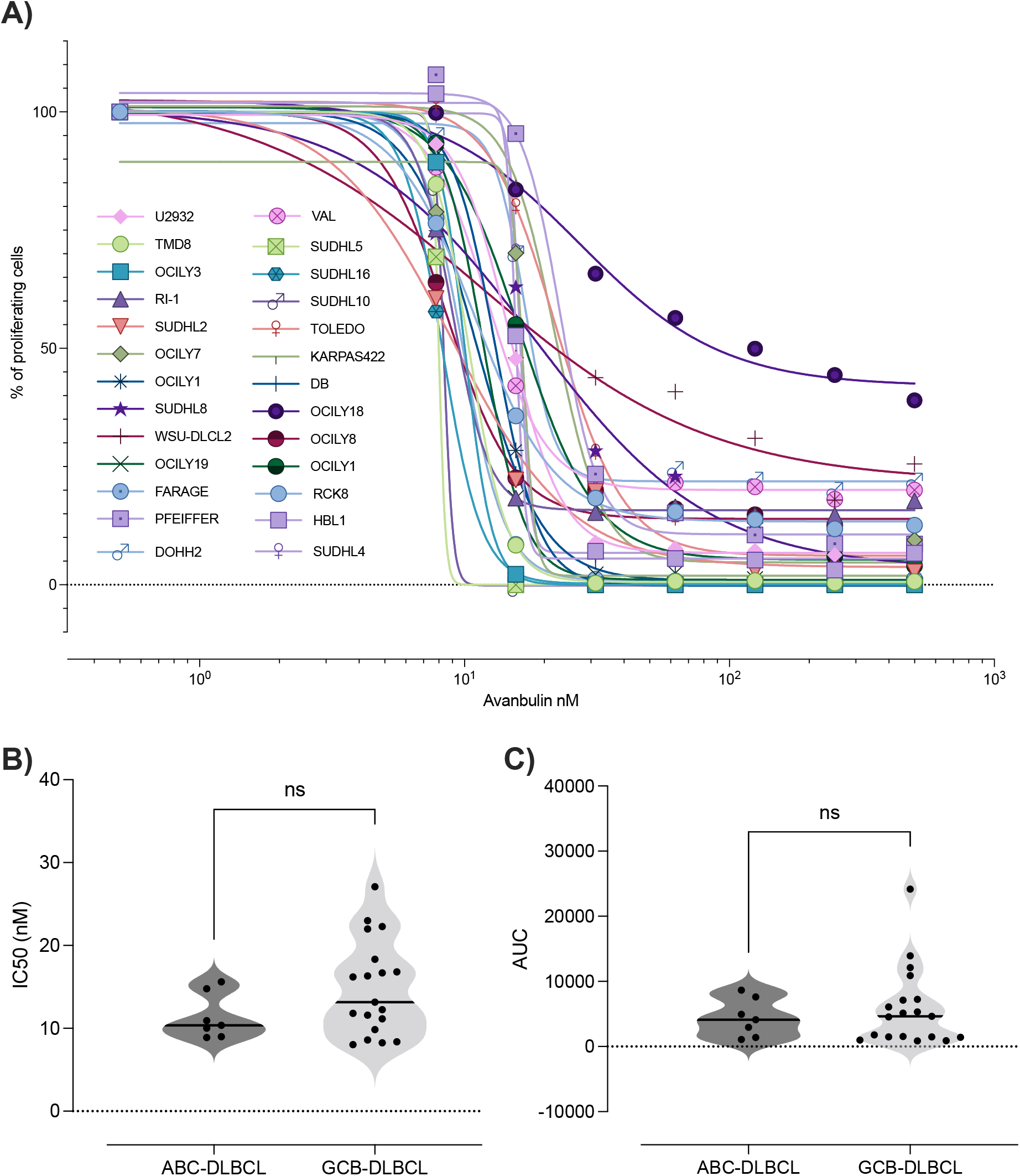
Anti-lymphoma activity of Avanbulin in diffuse large B cell lymphoma cell lines. A) Dose response curve of avanbulin in 26 human diffuse large B-cell lymphoma cell lines after 72 hours of treatment. B) IC50s and C) area under the curve comparison between ABC and GCB-DLBCL cell lines.

Cell growth and apoptosis were also monitored using a real-time quantitative live-cell imaging approach in a subset of DLBCL cell lines (n.=17, selected based on their capacity to grow in adherence), treated with DMSO or four different concentrations of avanbulin (5, 10, 20 and 40 nM) compared to untreated cells and images of cell lines were taken every 4 hours for a total of 72 hours. Cell numbers and apoptotic events were measured every 4 hours during a 72 hours period. Most of the cell lines showed a decrease in proliferation after treatment with BAL27862 in line with results obtained with MTT (Figure 2, Supplementary Table 2). Apoptosis induction was seen in 15 out of 17 cell lines tested at 20 and 40 nM, and only in one cell line already at 10 nM. Apoptosis could be detected in 8 of 15 cell lines already during the first 24 hours of treatment, while of the remaining cells 6 between 24 and 48 hours in six and after 48 hours in the remaining one (Supplementary Figure 1, Supplementary Table 2).

**Figure 2.**
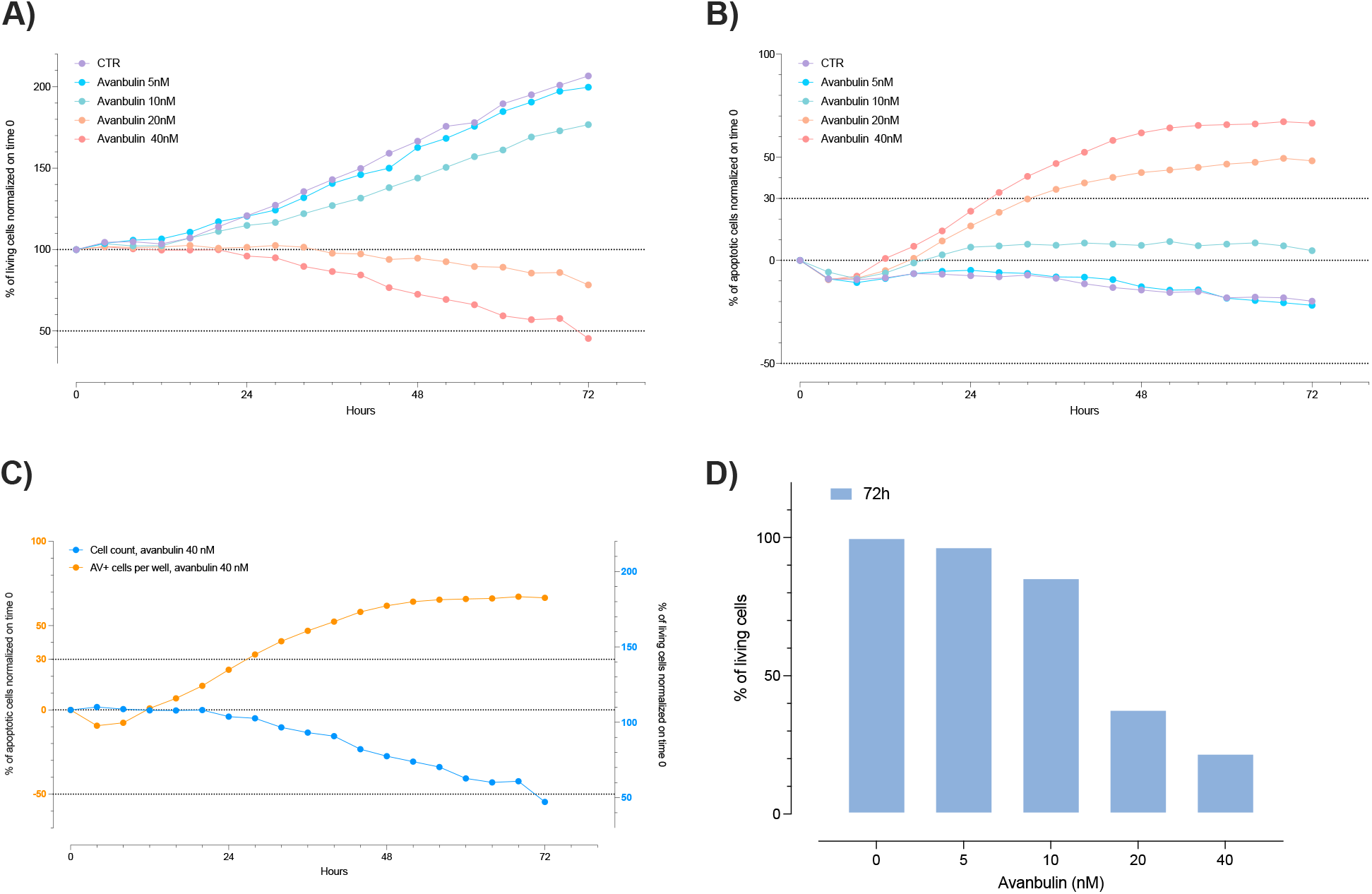
Antiproliferative activity and apoptosis induction of Avanbulin. Representative live cell imaging analysis performed in U2932 cell line. Cells under avanbulin treatment (5, 10, 20, 40 nM) compared to untreated cells A) Cell growth at different time points under avanbulin treatment. B) Apoptosis induction during treatment measured as percentage of apoptotic cells normalized at the beginning of the treatment. C) Comparison of cell growth and apoptosis induction at 40nM. D) Percentage of living cells at 72 hours normalized on untreated cells.

We did not observe an association between sensitivity and TP53 or BCL2 status, while based on AUC we identified an association between MYC translocation and sensitivity, with wild type cells more sensitive to the treatment (Supplementary Figure 2, Supplementary Table 3).

Cell cycle experiments were performed in two ABC (OCILY3 and TMD8) and 2 GCB (SUDHL5 and SUDHL16) DLBCL cells at 24, 48 and 72 hours. A strong cytotoxic effect of avanbulin was confirmed with a time-dependent accumulation of cells in sub-G0, starting already at 24 hours in all the four cell lines. A G2-M arrest was also seen in 3 out of 4 cell lines, more evident at 24 hours (Supplementary Figure 3).

### EB1 and MYC expression in cell lines

Based on the reported possible association between EB1 and sensitivity to avanbulin ^18–20^, we evaluated RNA expression in lymphoma cell lines, with no differences between GCB and ABC DLBCL (Supplementary Figure 4). All lymphoma cell lines expressed a high level of EB1. No correlation was observed between expression levels of EB1 and MYC (Supplementary Figure 4.), reported as positively regulated by EB1 in colorectal cancers ^21^. When we looked at the possible correlation between EB1 or MYC RNA and protein expression and avanbulin activity, we did not identify any trend for higher sensitivity in EB1 or MYC higher expressing cells comparing low expressors. Most of the cell lines were highly sensitive to avanbulin together with a general high EB1 and MYC expression (Supplementary Figure 4, Supplementary Table 4),

### EB1 and MYC expression in DLBCL clinical specimens

We then assessed EB1 RNA expression in DLBCL clinical specimens taking advantage of three publicly available gene expression datasets (GSE4475 ^25^, n.=221; GSE10846 ^29^, n.=420; GSE22470 ^26^, n=271). The transcript was highly expressed in most of the clinical specimens, with no difference based on the cell of origin (Figure 3.). In agreement with what was seen in cell lines, we did not observe any correlation between EB1 RNA expression and MYC RNA expression and no association between EB1 expression and the presence of *MYC* translocation (Figure 3).

**Figure 3.**
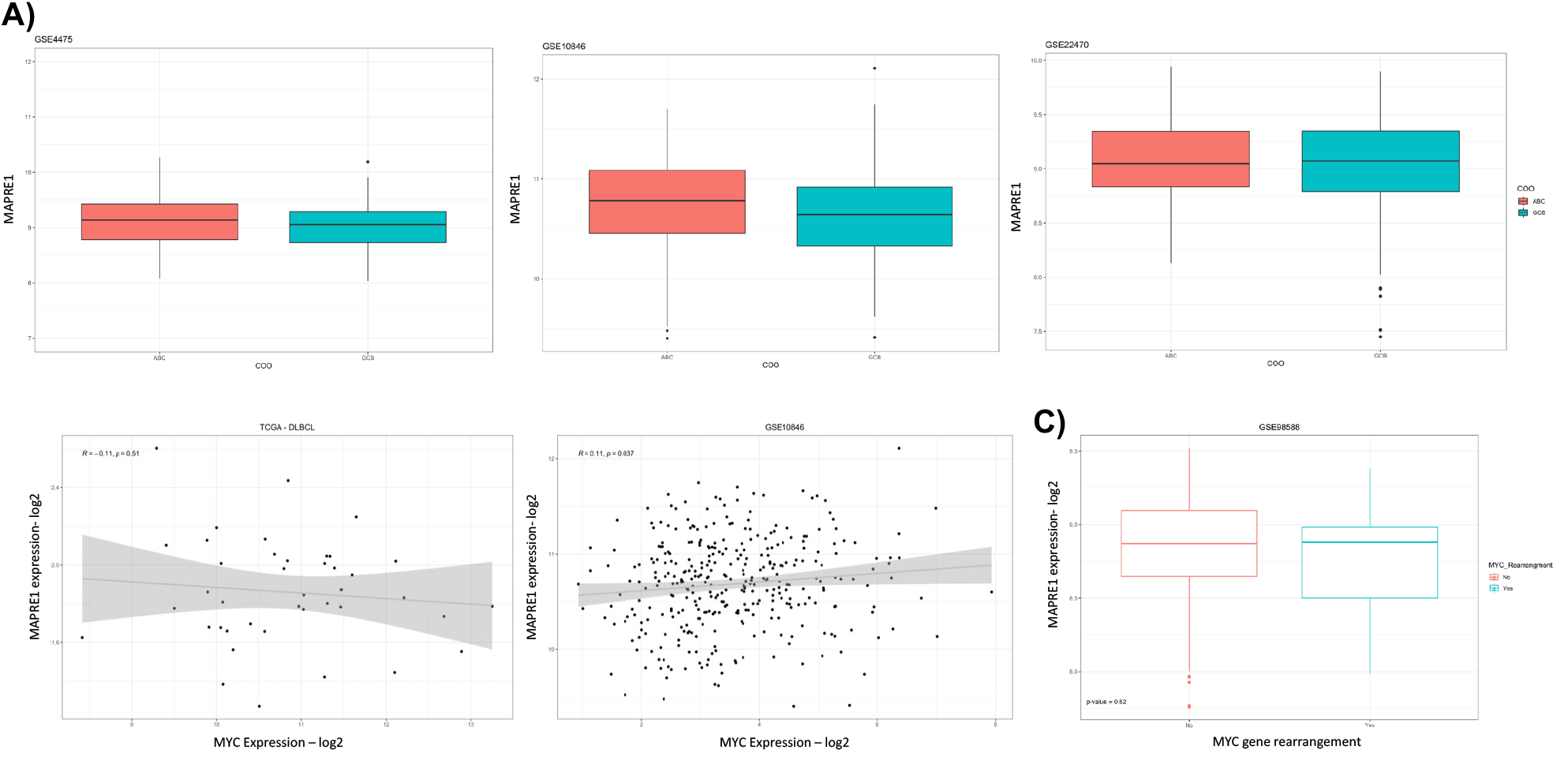
MAPRE-1 (EB1) RNA expression in DLBCL clinical specimens and correlation with MYC. A) MAPRE-1 (EB1) RNA expression in DLBCL clinical specimens divided by ABC (red) and GCB (blue) DLBCL. Three different datasets analyzed (GSE4475, GSE10846, GSE22470). B) Correlation between EB1 (MAPRE1 gene) RNA expression and MYC RNA expression in TCGA and GSE10846 datasets. C) MAPRE-1 RNA expression levels in DLBCL clinical specimen divided between MYC without (red) or with (blue) rearrangements.

Finally, we looked at EB1 protein expression by immunohistochemistry (IHC) in DLBCL clinical specimens. On a total of 101 cases analyzed on TMAs, 50 negative (49.5%) and 51 were positive (50.5%) (Supplementary Table 5). Among the latter, two percent of the tumor samples exhibited a moderate or strong EB1 staining in >50% of the tumor cells, 11% in 26-50% and 38% in 5-25%.

## Discussion

Here, we showed that the MTA avanbulin (BAL27862), the active moiety of lisavanbulin (BAL101553), has a potent *in vitro* anti-tumor activity in DLBCL, and that the microtubule associated protein EB1, a potentially predictive biomarker of sensitivity to the compounds, is highly expressed in both DLBCL cell lines and clinical specimens.

Avanbulin showed a median IC50 of 11 nM across 26 DLBCL cell lines after 72 hours of drug exposure. The activity was mostly cytotoxic as confirmed by both cell cycle analysis and apoptosis assays. Real-time quantitative live-cell imaging allowed us to observe a fast decrease in cell proliferation and apoptosis induction. Around half of the cell lines tested had apoptosis induction in the first 24 hours of treatment; the remaining half after one additional day of drug exposure. Apoptosis was preceded by G2/M arrest. Out of 17 cell lines tested 15 presented apoptosis induction already at 20 nM. These data are in line with what is reported in solid tumor models ^10–13^, although our results were not further *in vivo* validated.

The microtubule-associated protein EB1 has been indicated as a potential biomarker of sensitivity to avanbulin and lisavanbulin, especially in the context of glioblastoma, in which a small number of cases express high protein levels, and these could be the case that most benefit from treatment with the MTA ^18–20^. In our DLBCL series, we did not observe a clear correlation between EB1 expression and avanbulin anti-tumor activity, but only a trend for higher sensitivity in high EB1 expressor cell lines. Importantly, we detected high expression levels of EB1 at the RNA level in DLBCL cell lines and in DLBCL clinical specimens. Moreover, IHC indicated protein expression in 50% of DLBCL clinical specimens. Among the positive clinical specimens, 40% showed over 20% of the tumor cells highly positive with intensity of the staining between 2 and 3. Hence, based on this data, lisavanbulin might prove to be an active agent in this population of patients.

Avanbulin binds to the colchicine site of tubulin and causes a destabilization of microtubules ^9,12^. Although the effect of avanbulin on microtubules phenotypically differs from that caused by vinblastine ^9,12^, lymphoma cells are highly sensitive to microtubule destabilizers, including vinblastine itself, vincristine and MMAE. Our data suggest that DLBCL patients could potentially benefit from lisavanbulin treatment. Moreover, since the avanbulin tubulin binding site differs from both vinca alkaloids and MMAE and it does not appear to be a MDR1 substrate ^5,7,9,12,14^, lisavanbulin might be an alternative strategy also in patients that have developed resistance to these agents via tubulin mutations or MDR1 overexpression. A matter of concern for treating these patients is the potential overlapping toxicities, especially peripheral neuropathies, common to all MTAs ^5,8,16–18^, which might represent a limiting factor for using lisavanbulin in individuals already exposed to vincristine, vinblastine or antibody drug conjugates containing a MTA as payload.

In conclusion, avanbulin showed strong anti-tumor activity in DLBCL cell lines and expression of its putative biomarker EB1, at both RNA and protein level, was observed in clinical specimens.

## Acknowledgements

This project was partially supported by institutional research funds from Basilea Pharmaceutica International Ltd, Allschwil, Switzerland and partially supported by the Swiss Cancer Research grant KFS-4713-02-2019. Thanks to Targos (Discovery Life Sciences Biomarker Services GmbH) for performing the EB1 IHC analysis.

## Authorship Contributions

FS designed and performed experiments, analyzed results, prepared figures, cowrote the manuscript; LA, LC designed and performed experiments, analyzed results, and prepared figures; SN, AR, GR, AJA: performed experiments; NFG, FBa, ME, HL provided reagents, support and contributed to data interpretation; FBe provided conception and study design supervision and cowrote the manuscript; all authors read and edited the manuscript.

