## supplementary methods and figures for "Assessment of lisavanbulin (BAL101553) as an anti-lymphoma agent"

### Supplementary Figures

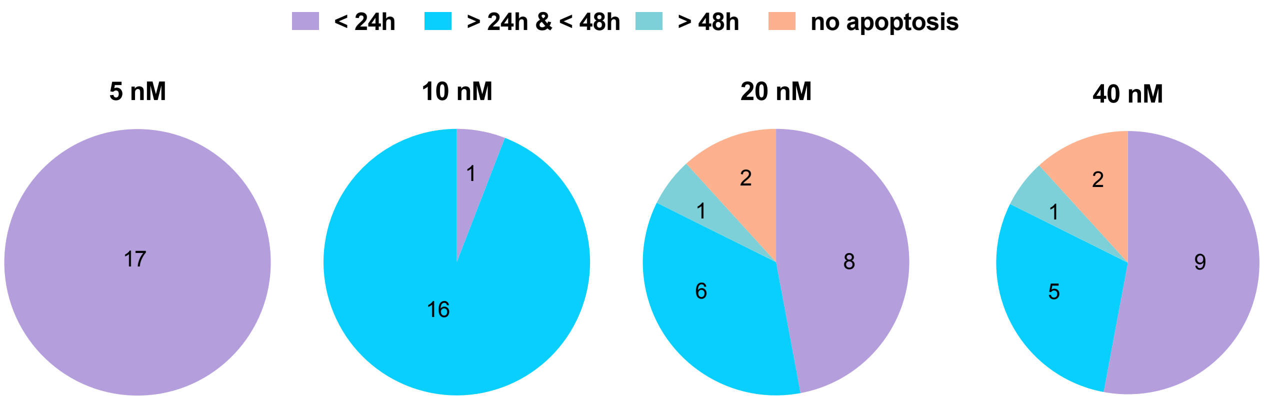


**Supplementary Figure 1. Number of cell lines with Annexin V induction**. Pie chart representing the number of cell lines undergoing apoptosis at different concentrations. Cells were divided based on the kinetics of apoptosis: no apoptosis, before 24, between 24 and 48 or later than 48 hours of treatment. Induction of apoptosis as >30% of apoptotic events compared to untreated cells by Incucyte.

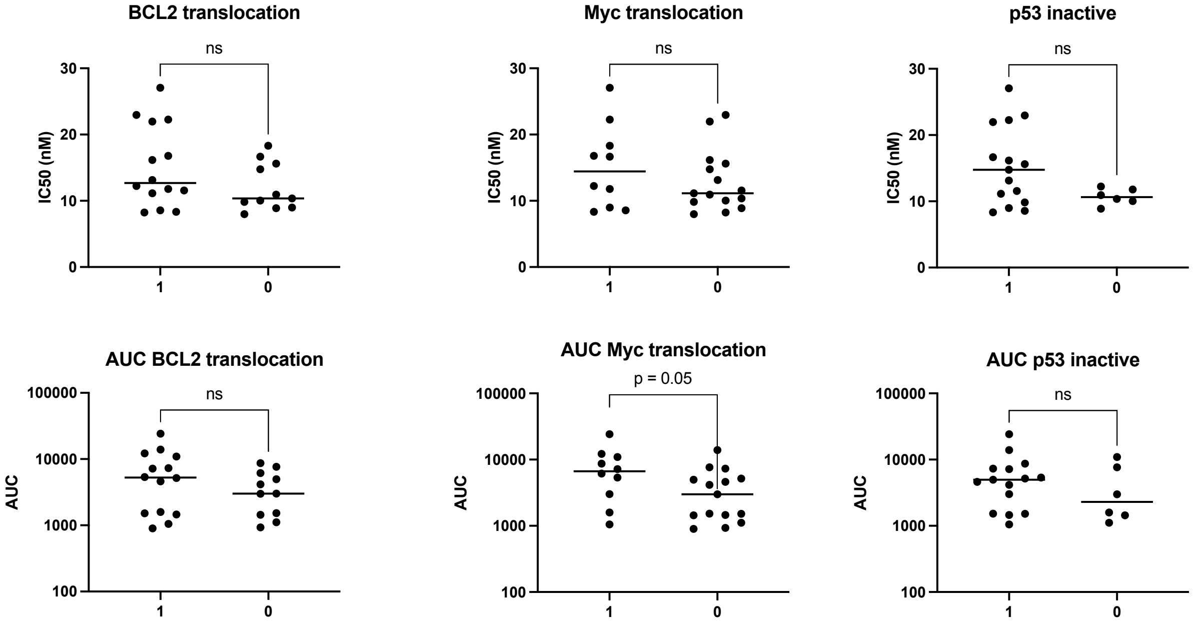


**Supplementary Figure 2. Association between sensitivity and MYC or BCL2 or TP53 status.** Association between IC50s or area under the curve (AUC) and MYC or BCL2 or TP53 status. 0 = wilde type; 1 = altered status (translocation or inactivation).

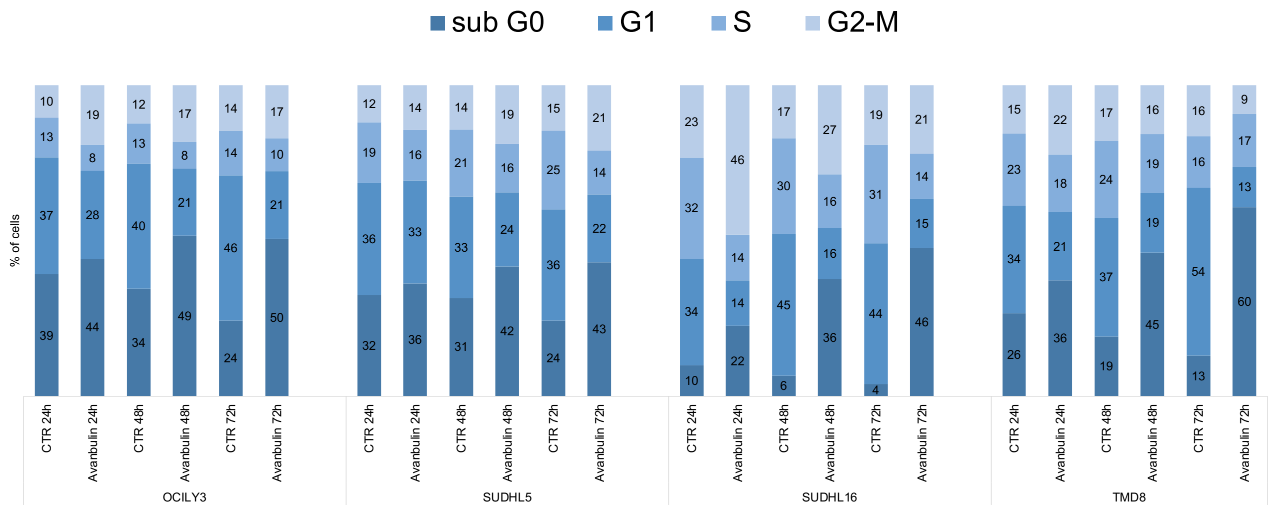


**Supplementary Figure 3. Cell cycle analysis.** Cell cycle analysis in four diffuse large b cell lymphoma cell lines after 24, 48 and 72 hours of treatment with 20nM of avanbulin. Average of at least two independent experiments.

**A)**

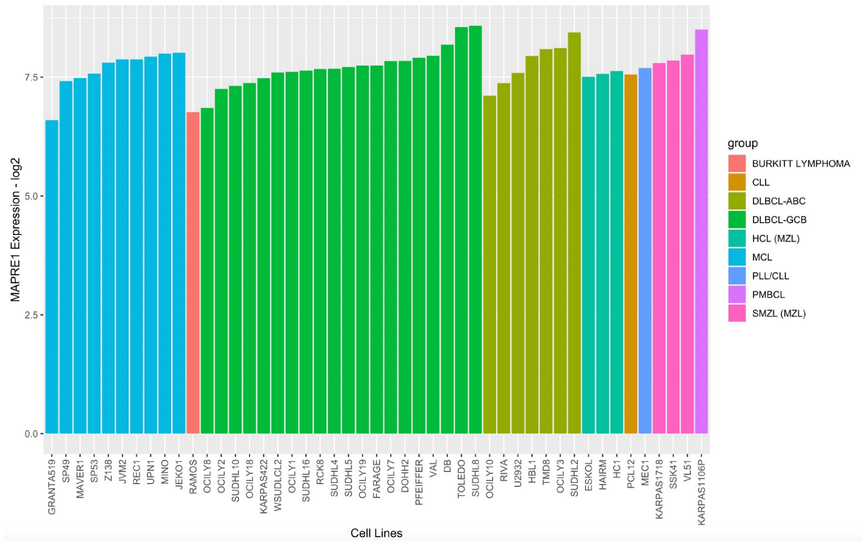


**B)**

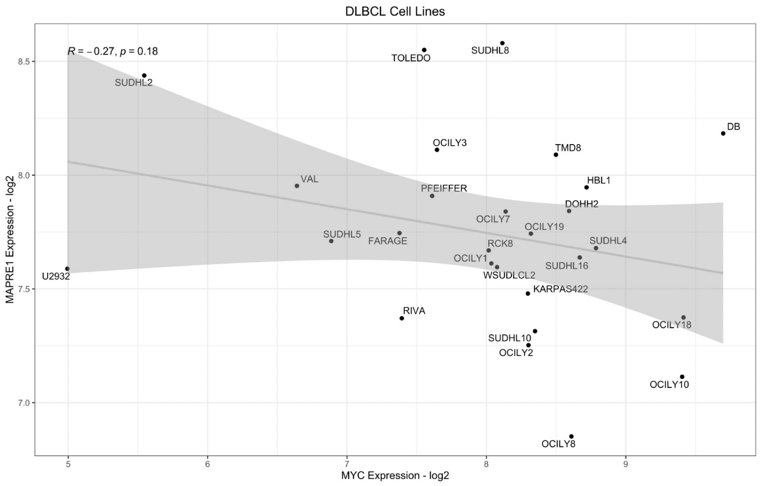


**Supplementary Figure 4. EB1 expression in lymphoma cell lines.**
A) EB1 (MAPRE1 gene) expression level lymphoma cell lines divided by subtype. B) Correlation between EB1 and MYC expression in DLBCL cell lines.

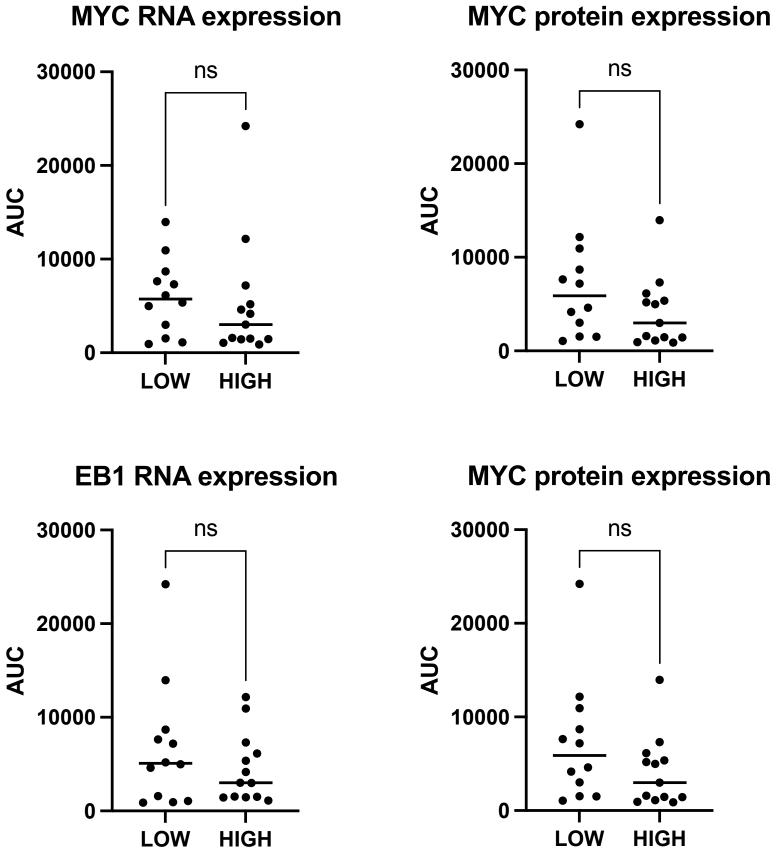


**Supplementary Figure 5. EB1 and MYC expression in sensitive and resistant DLBCL cell lines.** MYC and EB1 RNA or protein expression in cell lines with low (sensitive) or high (resistant) area under the curve (AUC). Cell lines were divided in low or high AUC if below or above the median AUC respectively.

**Supplementary Table 1. List of DLBCL cell lines with relative histology, IC50 and area under the curve (AUC) to lisavanbulin.**
Cell lines treated with increasing concentrations of lisavanbulin for 72 hours.

**Supplementary Table 2. Apoptosis induction by lisavanbulin in live cell imaging system (Incucyte).**
Cell lines treated with 4 different lisavanbulin concentrations (5, 10, 20 and 40 nM) and followed with cell imaging system for 72 hours.
Kinetics of annexin V induction represented as time (hour) when apoptosis starts.
Apoptosis induction defined as >30% of AV+ cells compared to untreated cells; 1 = apoptosis induction; 0 = no apoptosis induction.

**Supplementary Table 3. List of DLBCL cell lines with relative IC50 and AUC to lisavanbulin and MYC, BCL2 and p53 genetic alterations.**

**Supplementary Table 4. List of DLBCL cell lines with relative IC50 and AUC to lisavanbulin and MYC or EB1 (MAPRE1) gene and protein expression**

**Supplementary Table 5. Immunohistochemistry (IHC) data for EB1 staining in tissue microarrays of DLBCL patients.** LY1001D and LY2086B DLBCL patients tissue microarrays tested for EB1 protein expression. Intensity of EB1 expression was scored from 0 to 3. Cells with intensity 2 or 3 were considered positive and percentage of positive cells for each spot was reported.
